# Role of lipid nanodomains for inhibitory FcγRIIb function

**DOI:** 10.1101/2023.05.09.540011

**Authors:** Franziska Spiegel, Marius F.W. Trollmann, Sibel Kara, Matthias Pöhnl, Astrid F. Brandner, Falk Nimmerjahn, Anja Lux, Rainer A. Böckmann

## Abstract

The inhibitory Fcγ receptor FcγRIIb is involved in immune regulation and is known to localize to specific regions of the plasma membrane called lipid rafts. Previous studies suggested a link between the altered lateral receptor localization within the plasma membrane and the functional impairment of the FcγRIIb-I232T variant that is associated with systemic lupus erythematosus.

Here, we conducted microsecond all-atom molecular dynamics simulations and IgG binding assays to investigate the lipid nano-environment of FcγRIIb monomers and of the FcγRIIb-I232T mutant within a plasma membrane model, the orientation of the FcγRIIb ectodomain, and its accessibility to IgG ligands. In contrast to previously proposed models, our simulations indicated that FcγRIIb does not favor a cholesterol-or a sphingolipid-enriched lipid environment. Interestingly, cholesterol was depleted for all studied FcγRIIb variants within a 2-3 nm environment of the receptor, counteracting the usage of raft terminology for models on receptor functionality. Instead, the receptor interacts with lipids that have poly-unsaturated fatty acyl chains and with (poly-) anionic lipids within the cytosolic membrane leaflet.

We also found that FcγRIIb monomers adopt a conformation that is not suitable for binding to its IgG ligand, consistent with a lack of detectable binding of monomeric IgG in experiments on primary immune cells. However, our results propose that multivalent IgG complexes might stabilize FcγRIIb in a binding-competent conformation. We suggest differences in receptor complex formation within the membrane as a plausible cause of the altered membrane localization or clustering and the altered suppressive function of the FcγRIIb-I232T variant.

**Significance Statement:** Our study sheds new light on the molecular mechanisms underlying the regulation of immune signaling mediated by the inhibitory Fcγ receptor (FcγRIIb). By utilizing atomistic simulations and experimental assays, we demonstrate that FcγRIIb interacts with specific lipids in the plasma membrane. Notably, our findings challenge the current view of membrane heterogeneity in immune cells, as FcγRIIb is not localized in specialized membrane domains known as rafts. Rather, we propose that receptor complex formation modulates receptor localization and conformation, thereby enabling ligand binding.

Our findings have important implications for understanding how immune receptors function and communicate with each other, and may provide new opportunities for developing therapeutic strategies targeting FcγRIIb in diseases such as autoimmunity and cancer.

## Introduction

Fc receptors are single-pass transmembrane proteins, expressed on the surface of innate immune effector cells, that bind the fragment crystallizable (Fc) portion of antibodies. Immune effector cells such as monocytes, neutrophils, and macrophages are regulated by a precise interplay of activating and inhibitory receptors, which, upon ligand binding, contribute either a positive or a negative signal to the pro-inflammatory response [1]. Malfunctioning of these mechanisms can lead to excessive immune cell activation and the destruction of healthy tissue known as autoimmune pathology [2, 3]. One possible reason underlying an autoimmune response is the lack of negative signals, which are contributed by inhibitory receptors. In human, the only inhibitory Fcγ receptor (Fc receptors that bind specifically to immunoglobulin G, IgG, Fc portion), able to limit excessive immune cell activation through IgG is FcγRIIb [4], which is broadly expressed on innate immune effector cells and B cells. Activated by IgG binding to the extracellular domain of activating FcγRs in combination with FcγRIIb, the immunoreceptor tyrosine-based inhibitory motif (ITIM) located within the cytosolic domain of FcγRIIb is phosphorylated by Src kinases, resulting in the initiation of SHP or SHIP dependent inhibitory signaling pathways [5]. On B cells, which do not express activating FcγRs, FcγRIIb controls activating signals transduced via the B cell receptor (BCR). Loss of FcγRIIb function on B cells is consequently associated with an increased likelihood to produce self-reactive antibodies (autoantibodies), which may trigger autoimmune diseases such as systemic *lupus erythematosus* (SLE) [1, 3, 6–8].

Of note, a single nucleotide polymorphism (SNP) in FcγRIIb, changing an isoleucine to a thre-onine residue (I232T) in the sequence of the FcγRIIb transmembrane (TM) domain represents one of the strongest disease associations [9–13]. In addition, several pre-clinical studies have documented that humanized mice of this FcγRIIb genotype are more prone to produce self-reactive immune responses [14, 15]. Consequently, different studies aimed to explain the role of the I232T polymorphism in the TM domain of FcγRIIb for its inhibitory function [16].

According to Floto *et al*. [17] and Kono *et al*. [18], FcγRIIb-232T shows a decreased affinity to lipid rafts compared to the FcγRIIb-232I wild type (wt). As protein kinases phosphorylating the ITIM of FcγRIIb are preferentially located in lipid-raft domains, the exclusion of 232T from raft domains was suggested to weaken its inhibitory function [17–19]. Upon IgG binding, FcγRIIb downregulates the activation of the B cell receptor (BCR). One proposed mechanism for the inhibition of BCR signaling is the blocking of conformational changes in the cytoplasmic domains of the BCR that are normally involved in initiating signaling [20, 21]. Furthermore, FcγRIIb also hinders BCR oligomerization [20].

Xu and colleagues [23] suggested a different mechanism for the action of FcγRIIb and lack of action for the FcγRIIb-I232T mutant: They reported that the FcγRIIb TM domain may exert its function by blocking the colocalization of BCR and CD19. CD19 lowers the activation threshold of the BCR, i.e. the CD19-BCR colocalization renders IgG-induced BCR activation more likely. The I232T TM mutation fails to block this colocalization, thus suggesting a TM-based mechanism for the lack of activation control in SLE patients. Xu *et al*. [24] also state that I232T impairs the lateral mobility of FcγRIIb such that it diffuses too slowly to meet IgG at the same time as the BCR [24]. If the BCR recognizes IgG containing immune complexes (IC) at higher rates than FcγRIIb, the latter is very unlikely to inhibit BCR signaling, as it is already initiated by the time FcγRIIb arrives.

In 2019, Hu *et al*. [8] reported that the single nucleotide polymorphism from isoleucine to threonine leads to a kink within the TM helix of the disease-associated mutant, caused by the formation of an additional hydrogen bond between Thr^232^ and Val^228^, and to an increased tilt of the mutant TM domain with respect to the membrane. These changes result in a greater inclination of the FcγRIIb ectodomain towards the membrane, bringing the IgG binding site closer to the membrane. The proximity of the binding site and the membrane sterically hinder IgG from binding to the receptor, thereby leading to a loss of its suppressive function [8].

Altogether, previous studies reported a coupling between FcγRIIb function, receptor conformation, and receptor localization to specific membrane domains. Loss of function of the I232T mutant was attributed either to different domain localization of the TM mutant or to differences in membrane embedding and connected ligand accessibility suggesting membrane-related mechanisms underlying the disturbed allosteric regulation in patients suffering from SLE. Here, we scrutinize the membrane-receptor interaction and its influence on ligand binding (i) in multi-microsecond atomistic simulations of the wt receptor and of the I232T receptor mutant embedded in an asymmetric lipid bilayer model that relates to the recently reported composition of the plasma cell membrane of erythrocytes [25], and (ii) link our simulation studies to binding assays for IgG immune complexes (IC) to primary human B cells.

## Results

The membrane embedding characteristics of the FcγRIIb wild type receptor or its I232T mutant were systematically compared based on 20 μs long atomistic MD simulations. The different systems are listed in Table 1, the setups are further detailed in the Methods Section. For comparison, a glycosylated FcγRIIb was investigated as well [26]. Obtained results did not show significant differences to the unglycosylated variant (see SI). The extracellular leaflet of the plasma cell membrane model mimicking the composition of erythrocyte plasma membranes [25] is characterized by lipids with phosphatidylcholine (PC) headgroups and either glycerol or sphingosine (SM) backbones, and with mostly saturated or single unsaturated acyl chains (SI Table S1). The degree of unsaturation is significantly increased for lipids within the cytosolic leaflet. In addition to PC lipids, the cytosolic leaflet mainly comprises phosphatidylethanolamines (PE) and anionic phosphatidylserine (PS) lipids, as well as a few phosphatidylinositol (PI) lipids that frequently act as signaling lipids (charge 4 *e*^−^).

**Table 1:** Simulation systems. All systems are composed of a single FcγRIIb (residues 46-250) embedded in an asymmetric model membrane that mimics the composition of the plasma membrane [25], compare Table S1. The structure of FcγRIIb was combined from the crystal structure of the ectodomain (residues 46-217) [27] and the TM domain modeled in α-helical conformation.

| System | Description | Number of replicas | simulation length |
| --- | --- | --- | --- |
| $Fc\gamma RIIb^{wt}$ | $Fc\gamma RIIb$ wild type | 9 | 20 $\mu s$ |
| $Fc\gamma RIIb^{mut}$ | $Fc\gamma RIIb$ I232T mutant | 3 | 20 $\mu s$ |

The asymmetric plasma membrane model consists of in total 16 different lipid types (compare SI Table S1). Analysis of lipid diffusion shows that lipids within the extracellular leaflet move systematically slower than those located within the cytosolic leaflet (SI Figure S1). The averaged diffusion coefficients for the different leaflets of the FcγRIIb^*wt*^ system were *D* = (6.7 *±* 0.2) *·* 10^−9^ cm^2^/s for the extracellular leaflet and *D* = (23.8 *±* 0.5) *·* 10^−9^ cm^2^/s for the cytosolic leaflet, with similar coefficients for the different lipid types within the respective leaflet (similar values for FcγRIIb^*mut*^). I.e., lipid diffusion was faster by a factor of ≈ 3 within the cytosolic leaflet. During the 20 μs simulation time, lipids within the extracellular and cytosolic leaflets cover a mean distance of 7.3 nm and 13.8 nm, respectively.

### Receptor membrane embedding, receptor conformation, and IgG binding

All studied receptors displayed stability of the TM helix and of the ectodomain when embedded in the plasma membrane model. For all simulations, the root mean square deviations (*rmsd*) were at or below ≈2 Å for the TM helix. The receptor ectodomain is composed of two immunoglobulin(Ig)-like domains [27]. Despite the hydrogen-bonded anchoring of the two domains reported in the crystal structure [27], the domains are rather flexibly arranged. The *rmsd* of the Ig-like domains was typically between 3-4 Å. In one wt simulation (of in total 9 wt replica simulations), the Fc binding Ig-like domain partially unfolded upon membrane interaction.

The receptor TM helix tilt angle of ≈ 14^*°*^ (distribution width 6^*°*^) was unchanged between the wild type and the I232T mutant (Fig. 2 A). Noteworthy, previous simulation studies of FcγRIIb based on 200 ns simulations (analysis on final 80 ns) reported an increased tilt of the I232T mutant [8, 24]. This discrepancy is probably attributable to the 300 – 2,000 fold improved statistics here compared to previous studies (CHARMM36m force field [28–30] used here and in [8, 24]). The increased tilt was attributed to a kink within the transmembrane helix introduced by an additional hydrogen bond between threonine at position 232 and valine at position 228. Also here, our microsecond MD simulations suggest a different picture (Fig. 2 B): The TM domain kink angle was similar for all studied Fc receptors (mean of 164.0^*°*^ for wt, width of 8.7^*°*^). I.e., the additional hydrogen bond that may form between the hydroxyl group of Thr^232^ and the backbone carbonyl group of Val^228^ within the I232T mutant does not affect the TM orientation within the plasma membrane.

The difference in TM domain tilting of FcγRIIb and the I232T mutant observed in previous simulations led the authors to hypothesize that the change in tilt causes an increased bending of the ectodomain towards the plasma membrane for the mutant. The latter would likely be connected to a hindered IgG antibody binding to the Fc receptor and thereby explain the deleterious effect of the I232T mutation. In order to quantitatively characterize the ectodomain configurations sampled in our multi-microsecond MD simulations, we analyzed the distributions of inclination angles θ and ectodomain rotation angles γ that are summarized in Figures 2 D-F. Each sampled configuration of in total 280 μs simulation data was analyzed for possible binding of IgG avoiding sterical clashes in particular with the surrounding membrane. Regions compatible with IgG binding are shaded in gray. All simulations started from an upright oriented ectodomain configuration that is accessible to IgG (see Fig. 1 A).

**Figure 1:**
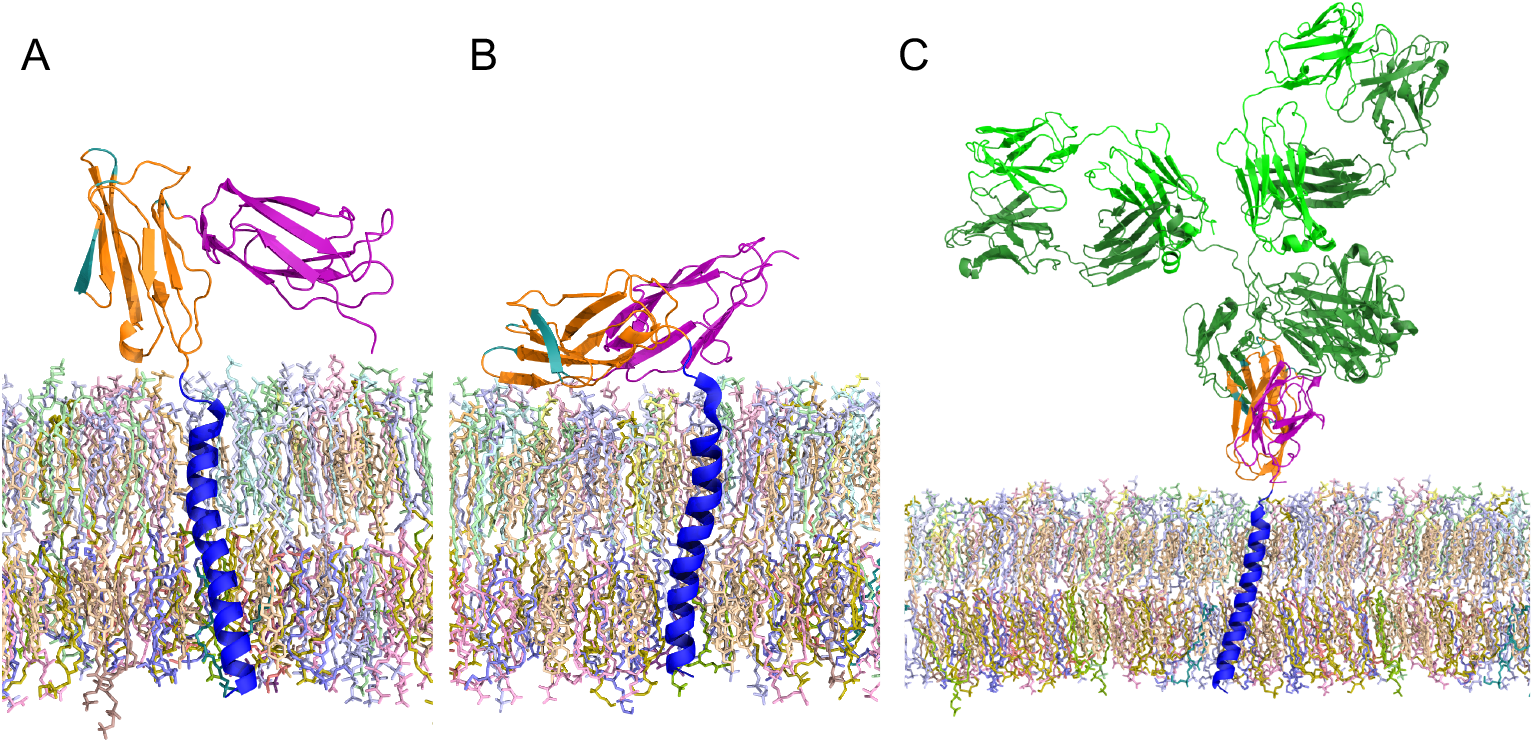
Snapshots of different FcγRIIb conformations. (*A*) Upright and (*B*) bent ectodomain configuration. The FcγRIIb TM domain is colored in *blue*, the ectodomain in *magenta* (Ig-like C2-type 1) and *orange* (Ig-like C2-type 2). The IgG binding site (defined as FcγRIIb residues within 5 Å of IgG residues) is highlighted in *teal*. (*C*) Structure of IgG (colored in *green*) bound to FcγRIIb, modeled after fit of the IgG-FcγR crystal structure [22] to a simulation snapshot of FcγRIIb (after 18 ns, ectodomain inclination angle of 11^*°*^). Lipids belonging to different lipid types are colored differently. For clarity, water and ions are not shown.

**Figure 2:**
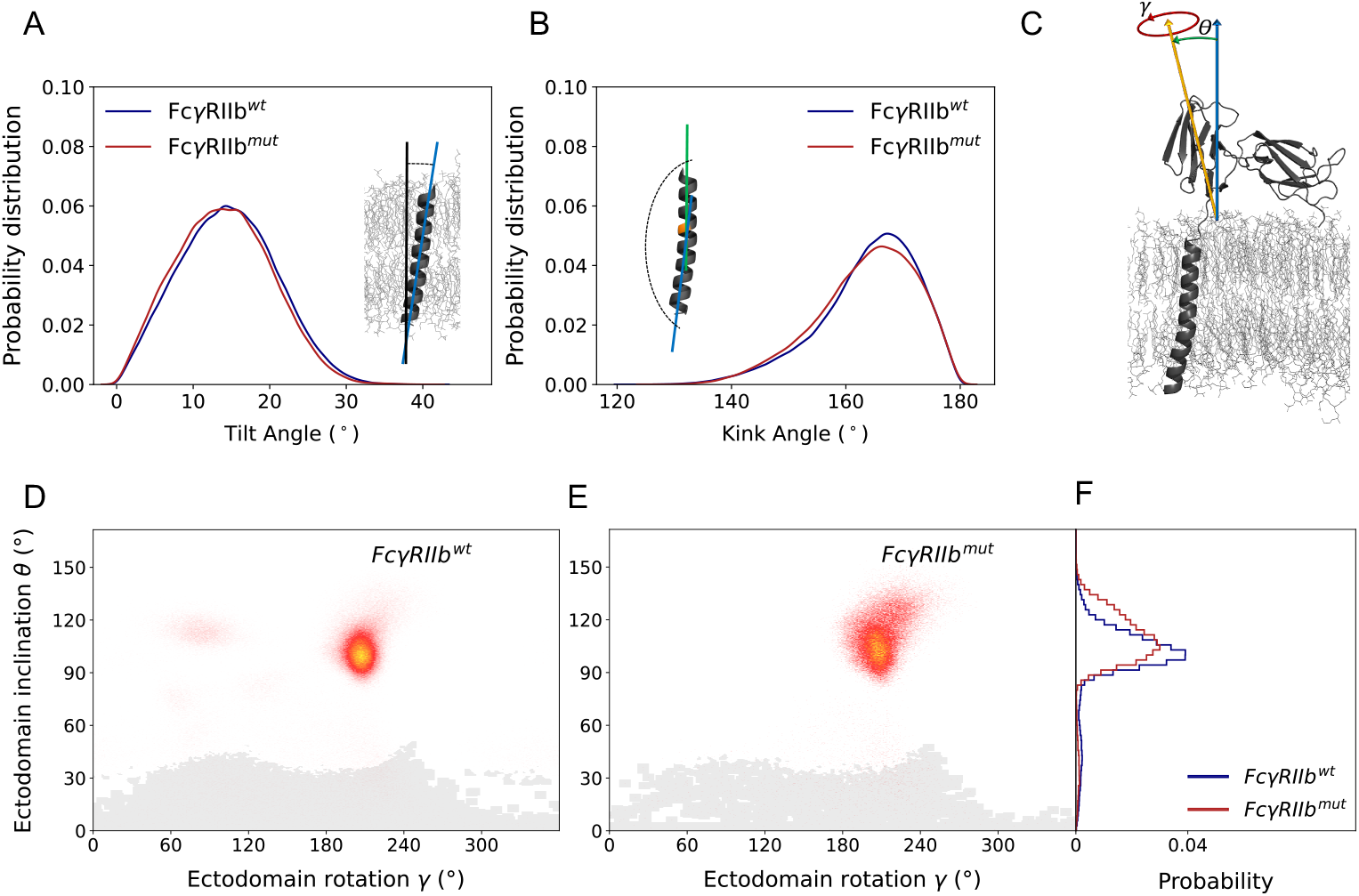
FcγRIIb transmembrane and ectodomain configurations sampled in multimicrosecond molecular dynamics simulations. (*A*) and (*B*) show the probability distributions of TM helix tilt and helix kink angles. (*C*) The ectodomain orientation with respect to the membrane is characterized using the ectodomain inclination angle θ and the ectodomain rotation angle γ. Panels (*D, E*) display the distribution of ectodomain configurations in the θ-γ space, including the data from all replicas (see Table 1), panel (*F*) the distribution of ectodomain inclination angles. FcγRIIb ectodomain orientations within the *gray shaded* area were compatible with the binding of IgG (4.4 % of all observed conformations for 232I, % for I232T.

As displayed in SI Fig. S2 for representative simulations, the ectodomain fluctuated in its orientation with respect to the membrane. After between 0.1 μs – 7 μs, the ectodomain bent towards the membrane in all wt and I232T mutant systems and stayed there, inclined, for the rest of the 20 μs simulations. No significant difference between the FcγRIIb wt and the I232T mutant were observed regarding the time until the inclination took place. While the mean inclination angle was 96.5^*°*^ for the wt, the I232T mutant showed a slightly increased inclination of 105.6^*°*^, compare Fig. 2 D,E). This difference, however, had no significant effect on the (sterically allowed) IgG binding ability which, in addition to the inclination angle, also takes the orientation of the IgG binding site with respect to the membrane surface into account. For strongly inclined conformations, i.e. for ectodomain configurations close to the membrane surface, no IgG binding was possible. Only very few of the overall sampled Fc ectodomain conformations were sterically compatible with a bound IgG (gray-shaded area in Fig. 2 D-E). Representative snapshots of IgG-accessible and non-accessible FcγRIIb conformations are displayed in Fig. 1.

### IgG immune complex binding in experiment

For validation of the simulation results, and in order to assess a potential functional impact of the FcγRIIb-I232T transmembrane polymorphism on binding of IgG, we compared IgG immune complex binding to primary human B cells. Due to the rare occurrence of the I232T transmembrane polymorphism in the human population we performed immune complex binding analysis for fresh peripheral blood samples (Figure 3 A,B) as well as frozen cord blood derived PBMC (SI Figures S6,S7). Employing an established IgG immune complex interaction assay [31], we quantified binding of all human IgG1-4 subclasses to isolated B cells with homozygous expression of either FcγRIIb-232I or -232T. In line with previous observations of IgG interaction with human FcγRIIb [31, 32], binding was most pronounced for IgG3 and IgG1 followed by IgG4 and IgG2. Most importantly, however, we did not observe significant differences in binding with respect to the two allelic variants of FcγRIIb (Figure 3 A). As receptor expression levels might have a crucial impact on IgG binding we confirmed comparable expression levels for both allelic receptor variants for a range of cord blood derived B cells (Figure 3 B), in line with previous reports studying fresh peripheral blood B cells [33].

**Figure 3:**
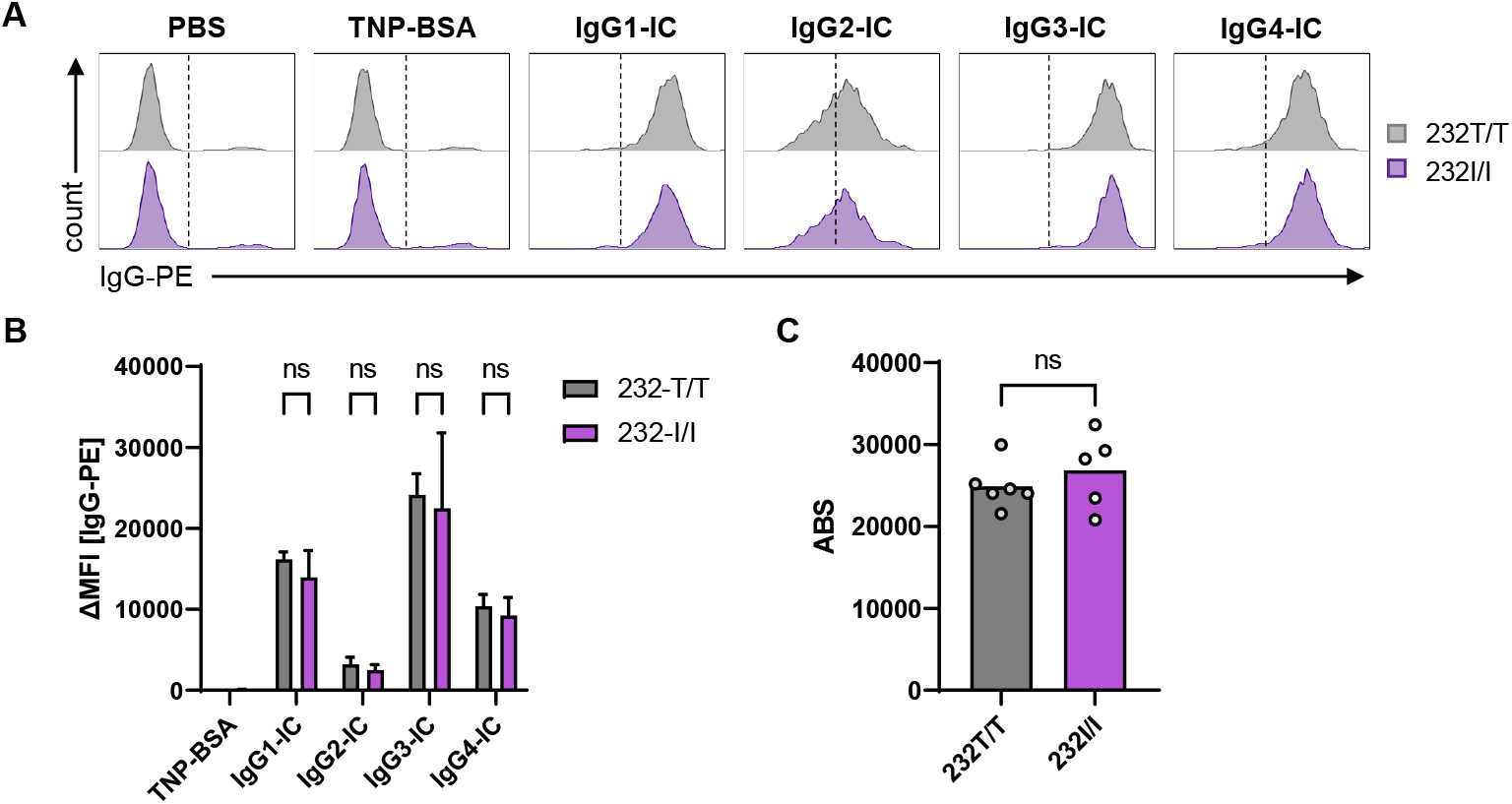
FcγRIIb-I232T polymorphism does not affect IgG-IC binding. Interaction of IgG1-4 immune complexes with primary B cells homozygously expressing either FcγRIIb-232I or -232T was assessed by flow cytometry. (*A*) Histogram display of one representative experiment shown. (*B*) Quantification of IgG-IC binding performed in technical replicates for three (232I, shown in *violet*) or two (232T, shown in *gray*) independent donors. Bars show mean and standard deviation of the median fluorescence intensity (MFI) upon detection of bound IgG-IC using fluorescently labeled anti-human IgG F(ab)2. TNP-BSA antigen is shown as negative control. (*C*) Quantification of FcγRIIb expression levels (depicted as antibody-binding sites, ABS) on cord blood derived B cells (*n* = 5 − 6 donors)

### Membrane analysis

Fc receptor activation has previously been discussed in the context of receptor partitioning to so-called raft domains [34]. The reported exclusion of the I232T mutant from rafts was suggested to contribute to the functional impairment of the mutant associated with SLE [17]. Thus, we characterized the diffusion and the nano-environment of FcγRIIb within a plasma membrane environment. We focused on specific receptor-lipid interactions, and the composition and order of the FcγRIIb receptor nano-environment.

### Receptor diffusion

The Fc receptor diffusion coefficient was *D* = (3.0 *±* 0.1) *·* 10^−9^ cm^2^/s for both FcγRIIb^*wt*^ and I232T mutant, in excellent agreement to the experimental values of *D*^*wt*^ = 3.3 *·* 10^−9^ cm^2^/s determined from 2D FRAP experiments for the wt receptor [24]. The slowed down diffusion of the mutant receptor with *D*^*mut*^ = 2.1 *·* 10^−9^ cm^2^/s reported in the latter experiments could thus not be explained based on the diffusion of single receptors in a biomembrane-like environment alone.

### Protein-lipid contacts

Specific lipid-protein interactions may act as a fingerprint for a particular protein [35]. Lipids either interact specifically with a receptor binding site or certain lipid species may be enriched in the vicinity of the protein due to the local lipid order or due to electrostatic interactions. Here, we assessed the lipid (number) density *ρ*_*N*_ (*r*) as a function of distance *r* from the receptor, normalized by the expected average density of the respective lipid (Figure 4). A value *ρ*_*N*_ (*r*_1_) = 1.5 thus corresponds to a mean lipid density within a distance of *r*_1_ from the receptor that is greater by 50% than the average density of the investigated lipid type within the simulation system.

**Figure 4:**
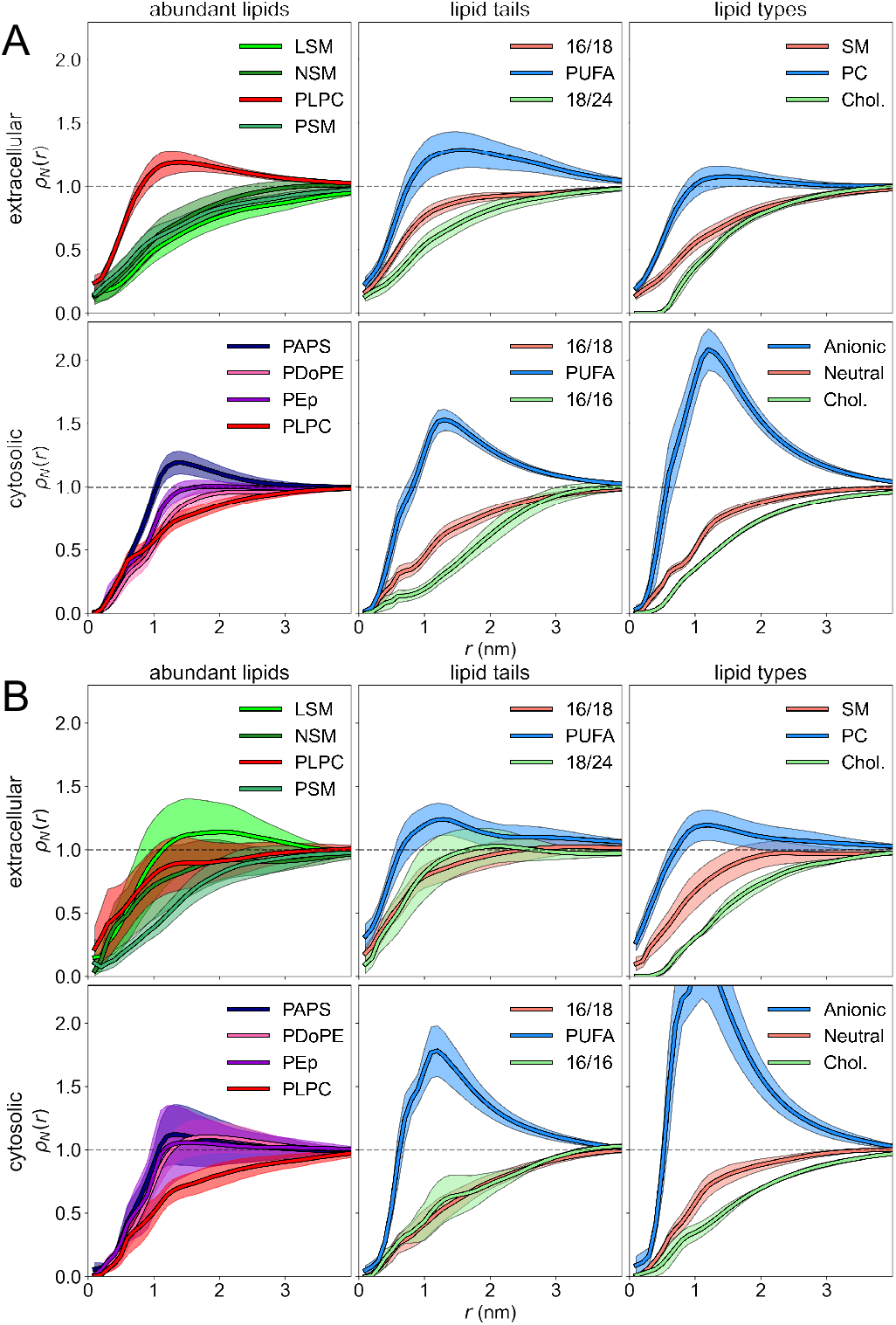
Lipid number density within extracellular and cytosolic leaflets of a plasma membrane model. The lipid densities *ρ*_*N*_ (*r*) are given as a function of distance *r* from the receptor, normalized by the expected average density of the respective lipid. The distances were computed to the center of mass of the leaflet’s matching apical region of the transmembrane helix for circular regions 0 ≤ *r*^*′*^ *< r*. Two datasets are shown, corresponding to the wt simulations (*A*) and I232T mutant simulations (*B*). Lipids were combined into different classes (see SI Table S1): Poly-unsaturated lipids (PUFA) include PAPS, PDoPE, SAPE, PIP2, SAPS, PAPC, PEp; 16/18 includes lipids with acyl chains of length 16 and 18 hydrocarbon groups (PLPC, PSM, SOPC, POPC, POPE); 16/16 is DPPC; 18/24 includes lipids with acyl chain lengths of 18 and 24 hydrocarbon groups (LSM, NSM); anionic lipids include PAPS, PIP2, and SAPS. Data were averaged over the final 10 μs of wt and I232T mutant simulations, with the shaded area representing the standard error of the mean calculated over all replica simulations.

Interestingly, for FcγRIIb, the density of lipids with phosphatidylcholine headgroups (PC) is overall increased in the extracellular leaflet within a distance of less than 2.5 nm from the receptor, the density of sphingomyelins and of cholesterol significantly decreased. Moreover, lipids with polyunsaturated fatty acyl chains (PUFAs) were preferred close to the receptor TM domain in both leaflets, in particular so for the cytosolic membrane leaflet. These observations are in contrast to the reported sorting of FcγRIIb to ordered, sphingomyelin and cholesterol-enriched raft domains.

Within the cytosolic leaflet, the density of the anionic phosphatidylserine (PS) lipids and of the highly negatively charged phosphatidylinositol-3,5-bisphosphate (PIP2) lipids are substantially enhanced close to the receptor. Figure 5 A shows the lateral positions of the PIP2 lipids within the cytosolic leaflet as a function of simulation time: For each simulation, 2-3 PIP2 lipids get tightly bound to the receptor (*filled gray circles* in Fig. 5 A), the PIP2 diffusion coefficient is substantially lowered by this complex formation (see SI Fig. S1). This binding is driven by a poly-basic stretch at the cytosolic part of the TM receptor domain. Different sampled FcγRIIb-PIP2 configurations are given in Figure 5 B-D.

**Figure 5:**
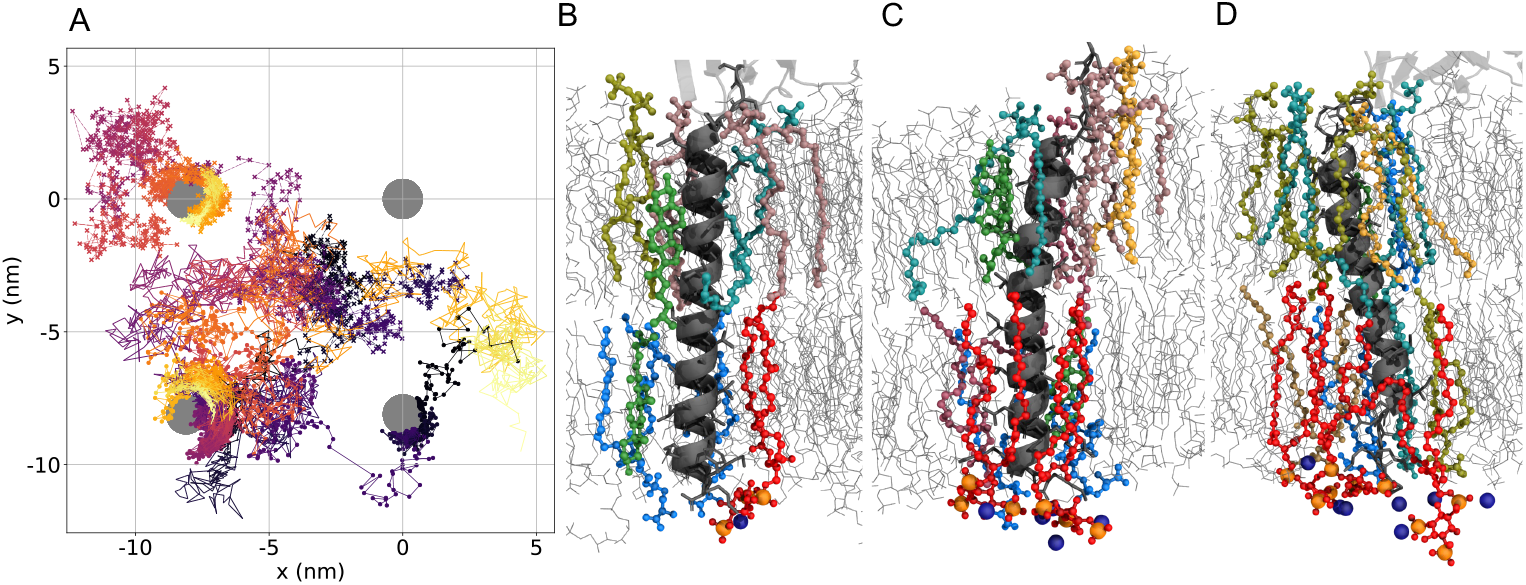
(*A*) Lateral position of PIP2 lipids for a simulation of FcγRIIb^*wt*^ embedded in a membrane model. The receptor position is depicted as a filled gray circle and replicated for four periodic images. PIP2 movements are shown as lines colored with gradient from dark purple to light yellow, tracking from *t* = 0μs to *t* = 20μs (see also SI Fig. S8 for PIP2 traces in other replicas). (*B*-*D*) Simulation snapshots of the lipid environment of the FcγRIIb TM helix. PIP2 lipids are colored in red with phosphate atoms highlighted in orange, PS in marine-blue, SM in teal, CHOL in lightgreen, PE in sand, PLPC in olive, PAPC in yellow, SOPC in rose and PEp in darkrose. Sodium ions within 5 Å of PIP2-atoms are shown as dark-blue spheres. For clarity, water and the remaining ions are omitted.

The lipid environment of the I232T Fc mutant (Fig. 4 B) resembles the one of the wild type receptor. Noteworthy, the density of sphingomyelins is *increased* with respect to the wild type receptor, as is the number of lipids with saturated or mono-unsaturated fatty acyl chains. Thereby, the mean lipid order close to the mutant receptor is slightly increased (see SI Fig. S9).

## Discussion

In previous studies, it was suggested that the FcγRIIb wild-type receptor is localized to lipid raft domains, while the SLE-associated FcγRIIb-I232T TM mutant receptor is observed to stay out-side of rafts, and a changed mutant receptor ectodomain conformation was related to the loss of function. These observations suggest a coupling between receptor localization and function.

This study conducted multi-microsecond all-atom molecular dynamics simulations to investigate the membrane embedment of FcγRIIb and of the FcγRIIb-I232T mutant within a plasma membrane model. The analysis focused in particular on the orientation of the FcγRIIb ectodomain and its accessibility to IgG ligands and the lipid nano-environment of this inhibitory Fc receptor.

Our simulation results reveal that all receptor variants exhibit a pronounced inclination of the receptor ectodomain towards the membrane surface, which is consistent across both the wild-type and the I232T mutant. As such, our findings suggest that the vast majority of FcγRIIb monomers, irrespective of the variant, adopt a conformation that is unsuitable for IgG interaction. These findings are consistent with previous reports of FcγRIIb^*wt*^, which indicate the absence or only a minimal binding of monomeric IgG both in cell-free systems [36] and on FcγRIIb-expressing cells [37].

Thus, we used IgG-immune complexes (IC) which represent the physiological ligand of low affinity FcγRs including FcγRIIb. In addition, we used IC binding to primary B cells to experimentally assess IgG binding to avoid artifacts due to altered receptor expression levels present in cultured cell lines. In fact, while FcγRIIb is also expressed on other human immune cell subsets, B cells are ideally suited because they lack expression of other activating FcγRs that might mask altered binding to FcγRIIb variants. Thus, IgG-IC binding to B cells can be solely attributed to FcγRIIb. Our data show no differences in IgG binding between B cells from homozygous FcγRIIb wildtype or mutant I232T donors, which is consistent with previous observations [19] but in contrast to the results published by Hu and colleagues [8]. Of note, we were able to detect binding of all IgG subclasses despite FcγRIIb being described as a non-binder for IgG2 due to a single nucleotide polymorphism [36]. This low affinity can be overcome by sufficiently multivalent IgG-IC [31, 38], as applied in our study, but not reflected by the experimental setup [39, 40] employed by Hu and colleagues [8].

However, we cannot exclude differing association or dissociation kinetics as flow cytometry experiments rather allow to quantify the level of equilibrium binding. Another limitation of the current study might be that binding was assessed at cold temperatures, which restricts lateral diffusion of membrane receptors, although previous studies have shown that mobility does not impact IgG binding by FcγRs [8, 40]. Our data suggest a model where multiple IgG Fc domains in multivalent immune complexes capture and stabilize FcγRIIb in a suitable conformation, which is not affected by the I232T polymorphism.

With regard to receptor domain localization, our simulation data show that FcγRIIb^*wt*^ monomers, as well as the I232T receptor mutant, are *not* located in domains enriched in cholesterol or sphingolipids, referred to as raft domains. In contrast, we found that cholesterol was depleted for all studied FcγRIIb variants within a 2-3 nm environment of the receptor. Raft domains are further characterized by an increased lipid order and thus reduced diffusional dynamics. At variance with this raft perspective, a *reduced* lateral mobility of the non-raft FcγRIIb-I232T mutant was reported from single-molecule imaging [24], i.e. a reduced dynamics of the receptor variant within a less ordered and likely more fluid environment. We suggest that this change in mobility is possibly related to a mutation-induced modified receptor clustering and not connected to a kinking of the TM domain and a related change in ectodomain configuration and ligand accessibility. Rather, a change in clustering may entail a reduced ligand accessibility. We note that we here used the asymmetric membrane lipid composition reported for red blood cells [25]. Thus, we cannot exclude effects on receptor nanodomains that would originate from different lipid compositions for different primary immune cell subtypes [41].

Instead of a raft environment, lipids with poly-unsaturated fatty acyl chains are enriched within the receptor nano-environment, harboring preferentially a phosphatidyl-choline (PC) headgroup in the extracellular leaflet and (poly-)anionic headgroups in the cytosolic leaflet of the plasma membrane.

We propose that the seemingly conflicting results for receptor localization, IgG-binding competent receptor conformation, and receptor mobility may be resolved by a yet unidentified receptor complex formation that is altered by the I232T TM mutation, as was recently described for monocytes, where the binding of Dectin-1 to FcγRIIb was shown to modulate FcγRIIb conformation leading to enhanced IgG binding [37]. Based on our MD simulations and antibody binding assays, we suggest that differences in FcγRIIb-related complex formation caused by the mutation are a likely cause for the reported differences in plasma membrane localization and IgG binding, as earlier suggested by Floto and colleagues [17]. Additionally, the preferential binding of the signaling lipid PIP2 to the cytosolic part of the receptor TM domain further suggests a possibly PIP2-driven receptor sequestering driven by electrostatic interactions, similar to the sequestration previously observed for the fusion peptide syntaxin-1A in neuronal cells [42].

Importantly, the lack of any raft characteristics within the FcγRIIb nanoenvironment suggests an alternative view on plasma membrane heterogeneity [43]: There are likely no underlying unique nano-or microdomains that function to compartmentalize or cluster immune signaling. Rather, receptors likely shape their specific lipid environment i.e. a specific immune domain that may contribute to receptor complex formation, clustering, and immune signaling.

## Materials and Methods

### MD simulations

See SI Appendix for a detailed description of the simulation setup and analysis.

### Isolation of genomic DNA and genotyping of FcγRIIb-232I/T

Genotyping of blood samples for FcγRIIb alleles was done as described before [14]. Briefly, genomic DNA was isolated with the „QiaAmp DSP Blood Mini Kit “(Qiagen, Hilden) according to the manufacturer’s instructions. For FcγRIIb genotyping a two-step PCR protocol was followed: Firstly, a 15 kb product was amplified using the Qiagen ‘LongRange PCR Kit’ (Qiagen) (Primers LongRange fwd: ctccacaggttactcgtttctaccttatcttac and LongRange rev: gcttgcgtggc-ccctggttctca) generating a 14.7 kb amplicon. Following purification by gel electrophoresis with the Qiagen ‘Gel purification Kit’, the PCR product was used as a template for the nested PCR (Primers R2Btm-fwd: aaggggagcccttccctctgtt and R2Btm-rev: gtggcccctggttctcagaa). Finally, the PCR product (2365 bp) containing the transmembrane region was purified and sequenced (using the R2Bseq primer: aaggggagcccttccctctgtt).

### IgG-IC binding assay

Peripheral blood mononuclear cells (PBMC) were isolated from peripheral blood or umbilical cord blood by density gradient centrifugation. 200,000 PBMCs were seeded into 96-well V-bottom plate. IgG immune complexes (IgG-IC) were generated as described before [31]. Briefly, 10 *μ*g/ml trinitrophenyl (TNP)-specific human IgG1-4 (clone 7B4) were incubated with 5 *μ*g/ml TNP-conjugated BSA (TNP-33-BSA, BioSearch Technologies) in PBS for 3 hours gently shaking at 60 rpm. PBMCs were incubated with IgG-IC for 1 hour on ice gently shaking. Unbound IgG-IC were removed by washing with PBS. Subsequently, cells were stained with fluorescently labelled antibodies specific for CD56 (clone MEM188, FITC-conjugated, Biolegend), CD3 (clone UCHT1, PerCP-conjugated, Biolegend), CD33 (clone WM53, APC -conjugated, Biolegend), CD19 (clone SJ25C1, PE/Cy7-conjugated, Biolegend), CD45 (clone 2D1, APC-Fire BV750-conjugated, Biolegend) and CD14 (clone M5E1, BV510-conjugated, Biolegend) for identification of PBMC subsets. For quantification of bound IgG-IC, cells were in addition stained with PE-conjugated goat anti-human IgG Fc F(ab)2 (Jackson Immunoresearch). Binding of TNP-33-BSA served as control, background fluorescence intensity of cells incubated with PBS was subtracted. Dead cells were excluded by staining with DAPI. Samples were acquired on a FACS CantoII (BD BioSciences) and analysed with BD FACSDiva and Prism software. Data sets were statistically compared using 2-way ANOVA.

### Quantification of FcγRIIb expression

FcγRIIb on B cells was quantified by measuring the Antibody Binding Capacity (ABC) for the FcγRIIb specific antibody (clone 2B6, PE-conjugated, produced in house). The ABC values were calculated using specific reference curves for the correlation between fluorescence intensity of a cell upon binding by anti-FcγRIIb-PE and the number of antibody binding sites. Reference curves were generated in each experiment using sets of Quantum Simply Cellular (QSC) microspheres (Bangs Laboratories Ltd.) with known numbers of antibody binding sites. Beads and cells were stained with the same antibody concentration [33]. Samples were acquired on a FACS CantoII (BD BioSciences) and analysed with BD FACSDiva and Prism software. Data sets were statistically compared using unpaired t-test following confirmation of Gaussian distribution.

## Data Archival

Raw simulation data will be deposited in the Zenodo repository after acceptance. All other data are included in the article and the SI Appendix.

## Acknowledgements

We thank Heike Albert and Prabha Varghese for expert technical assistance. The authors gratefully acknowledge the computer resources and support provided by the Erlangen Regional Computing Center (RRZE) and the Erlangen National High Performance Computing Center (NHR@FAU). The study was funded through grants from the German Research Foundation (NI711, TRR305-B02, CRC1526-A07 to FN) and the NIH (NIH/U01-AI148119-02 to FN).

## References

1. Nimmerjahn, F. & Ravetch, J. Fcγ receptors as regulators of immune responses. Nat. Rev. Immunol. 8, 34–47 (2008).

2. Sármay, G. Biologia Futura: Emerging antigen-specific therapies for autoimmune diseases. Biol Futura, 1–10 (2021).

3. Ben Mkaddem, S., Benhamou, M. & Monteiro, R. Understanding Fc receptor involvement in inflammatory diseases: from mechanisms to new therapeutic tools. Front. Immunol. 10, 811 (2019).

4. Takai, T., Ono, M., Hikida, M., Ohmori, H. & Ravetch, J. V. Augmented humoral and anaphylactic responses in Fc gamma RII-deficient mice. Nature 379, 346–349 (1996).

5. Bolland, S. & Ravetch, J. V. Inhibitory pathways triggered by ITIM-containing receptors. Adv. Immunol. 72, 149–177 (1999).

6. Niederer, H., Clatworthy, M., Willcocks, L. & Smith, K. FcγRIIB. FcγRIIIB, and systemic lupus erythematosus. Ann. N. Y. Acad. of Sci 1183, 69–88 (2010).

7. Hargreaves, C. et al. Fcγ receptors: genetic variation, function, and disease. Immunol. Rev. 268, 6–24 (2015).

8. Hu, W. et al. FcγRIIB-I232T polymorphic change allosterically suppresses ligand binding. Elife 8, e46689 (2019).

9. Willcocks, L. et al. A defunctioning polymorphism in FCGR2B is associated with protection against malaria but susceptibility to systemic lupus erythematosus. Proc. Natl. Acad. Sci. USA 107, 7881–7885 (2010).

10. Verbeek, J., Hirose, S. & Nishimura, H. The complex association of FcγRIIb with autoimmune susceptibility. Front. Immunol. 10, 2061 (2019).

11. Kyogoku, C. et al. Fcγ receptor gene polymorphisms in Japanese patients with systemic lupus erythematosus: contribution of FCGR2B to genetic susceptibility. Arthritis Rheumatol 46, 1242–1254 (2002).

12. Siriboonrit, U. et al. Association of Fcγ receptor IIb and IIIb polymorphisms with susceptibility to systemic lupus erythematosus in Thais. Tissue Antigens 61, 374–383 (2003).

13. Chu, Z. et al. Association of Fcγ receptor IIb polymorphism with susceptibility to systemic lupus erythematosus in Chinese: a common susceptibility gene in the Asian populations. Tissue Antigens 63, 21–27 (2004).

14. Baerenwaldt, A. et al. Fcγ receptor IIB (FcγRIIB) maintains humoral tolerance in the human immune system in vivo. Proc. Natl. Acad. Sci. USA 108, 18772–18777 (2011).

15. Danzer, H. et al. Human Fcγ-receptor IIb modulates pathogen-specific versus self-reactive antibody responses in lyme arthritis. Elife 9, e55319 (2020).

16. Wang, J., Li, Z., Xu, L., Yang, H. & Liu, W. Transmembrane domain dependent inhibitory function of FcγRIIB. Protein Cell 9, 1004–1012 (2018).

17. Floto, R. et al. Loss of function of a lupus-associated FcγRIIb polymorphism through exclusion from lipid rafts. Nat Med 11, 1056–1058 (2005).

18. Kono, H. et al. Spatial raft coalescence represents an initial step in FcγR signaling. J Immunol 169, 193–203 (2002).

19. Kono, H. et al. FcγRIIB Ile232Thr transmembrane polymorphism associated with human systemic lupus erythematosus decreases affinity to lipid rafts and attenuates inhibitory effects on B cell receptor signaling. Hum Mol Genet 14, 2881–2892 (2005).

20. Liu, W., Sohn, H., Tolar, P., Meckel, T. & Pierce, S. Antigen-induced oligomerization of the B cell receptor is an early target of FcγRIIB inhibition. J Immunol 184, 1977–1989 (2010).

21. Tolar, P., Sohn, H. & Pierce, S. The initiation of antigen-induced B cell antigen receptor signaling viewed in living cells by fluorescence resonance energy transfer. Nat Immunol 6, 1168–1176 (2005).

22. Falconer, D., Subedi, G., Marcella, A. & Barb, A. Antibody fucosylation lowers the FcγRIIIa/CD16a affinity by limiting the conformations sampled by the N162-glycan. ACS Chem. Biol 13, 2179–2189 (2018).

23. Xu, L. et al. Through an ITIM-independent mechanism the FcγRIIB blocks B cell activation by disrupting the colocalized microclustering of the B cell receptor and CD19. J. Immunol. 192, 5179–5191 (2014).

24. Xu, L. et al. Impairment on the lateral mobility induced by structural changes underlies the functional deficiency of the lupus-associated polymorphism FcγRIIB-T232. J. Exp. Med. 213, 2707–2727 (2016).

25. Lorent, J. et al. Plasma membranes are asymmetric in lipid unsaturation, packing and protein shape. Nat. Chem. Biol 16, 644–652 (2020).

26. Hayes, J. M. et al. Identification of Fc Gamma Receptor Glycoforms That Produce Differential Binding Kinetics for Rituximab. Mol. Cell. Proteomics 16, 1770–1788 (2017).

27. Sondermann, P., Huber, R. & Jacob, U. Crystal structure of the soluble form of the human Fcgamma-receptor IIb: a new member of the immunoglobulin superfamily at 1.7 A resolution. EMBO J. 18, 1095–1103 (1999).

28. Klauda, J. B. et al. Update of the CHARMM all-atom additive force field for lipids: validation on six lipid types. J. Phys. Chem. B 114, 7830–7843 (2010).

29. Huang, J. et al. CHARMM36m: an improved force field for folded and intrinsically disordered proteins. Nat. Methods 14, 71–73 (2017).

30. Yu, Y. & Klauda, J. B. Update of the CHARMM36 United Atom Chain Model for Hydrocarbons and Phospholipids. J. Phys. Chem. B 124, 6797–6812 (2020).

31. Lux, A., Yu, X., Scanlan, C. N. & Nimmerjahn, F. Impact of immune complex size and glycosylation on IgG binding to human FcγRs. J. Immunol. 190, 4315–4323 (2013).

32. Bruhns, P. Properties of mouse and human IgG receptors and their contribution to disease models. Blood 119, 5640–5649 (2012).

33. Kerntke, C., Nimmerjahn, F. & Biburger, M. There Is (Scientific) Strength in Numbers: A Comprehensive Quantitation of Fc Gamma Receptor Numbers on Human and Murine Peripheral Blood Leukocytes. Front. Immunol. 11, 118 (2020).

34. Pike, L. J. Rafts defined: A report on the Keystone symposium on lipid rafts and cell function. J. Lipid Res. 47, 1597–1598 (2006).

35. Corradi, V. et al. Lipid-Protein Interactions Are Unique Fingerprints for Membrane Proteins. ACS Cent. Sci. 4, 709–717 (2018).

36. Bruhns, P. et al. Specificity and affinity of human Fcgamma receptors and their polymorphic variants for human IgG subclasses. Blood 113, 3716–3725 (2009).

37. Seeling, M. et al. Immunoglobulin G-dependent inhibition of inflammatory bone remodeling requires pattern recognition receptor Dectin-1. Immunity (2023).

38. Robinett, R. A. et al. Dissecting FcγR Regulation through a Multivalent Binding Model. Cell Syst. 7, 41–48 (2018).

39. Chesla, S. E., Selvaraj, P. & Zhu, C. Measuring two-dimensional receptor-ligand binding kinetics by micropipette. Biophys. J. 75, 1553–1572 (1998).

40. Chesla, S. E., Li, P., Nagarajan, S., Selvaraj, P. & Zhu, C. The membrane anchor influences ligand binding two-dimensional kinetic rates and three-dimensional affinity of FcgammaRIII (CD16). J. Biomol. Chem. 275, 10235–10246 (2000).

41. Lühr, J. J. et al. Maturation of Monocyte-Derived DCs Leads to Increased Cellular Stiffness, Higher Membrane Fluidity, and Changed Lipid Composition. Front. Immunol. 11, 590121 (2020).

42. Van Den Bogaart, G. et al. Membrane protein sequestering by ionic protein-lipid interactions. Nature 479, 552–555 (2011).

43. Sevcsik, E. & Schütz, G. J. With or without rafts? Alternative views on cell membranes. Bioessays 38, 129– 139 (2016).

