## Supplementary Information for "Role of lipid nanodomains for inhibitory FcγRIIb function"

### Role of lipid nanodomains for inhibitory Fc$\gamma$RIIb function

**This PDF file includes:**

Supplementary text
Figs. S1 to S9
Table S1
References for SI reference citations

### Supporting Information Text

**Simulation systems.** The structure model of the human Fc$\gamma$RIIb including ectodomain and transmembrane (TM) domain (residues A46-K250) was combined from the crystal structure of the ectodomain (PDB code 2FCB (1), residues A46-P217) and the TM domain in $\alpha$-helical conformation (using the MODELLER software (2)). For the mutant receptor (I232T), the isoleucine at position 232 was replaced by threonine. N- and C-termini were capped with acetyl and amide groups, respectively. For comparison, a glycosylated Fc$\gamma$RIIb receptor was studied as well (system Fc$\gamma$RIIb$^{glyc}$). Studies on Fc$\gamma$R glycosylations indicated a large heterogeneity in their N-glycans with varying complexity but showed a common core motif consisting of D-Mannose (Man), N-Acetyl-D-glucosamine (GlcNAc), and 6-Deoxy-L-galactose (Fucose) (3–5). In the present work, N-glycans were modeled with a fucosylated Man3GlcNAc2Asn core sequence adopting a biantennary structure with one GlcNAc residue each (see Fig. S5). Three potential N-glycosylation sites were assigned by the Uniprot database (6) (ID: P31994, last accessed: 02.04.2023) that show the characteristic Asn-X-Ser/Thr motif for N-glycans (7).

A plasma membrane-like, asymmetrically composed lipid bilayer was constructed using the CHARMM-GUI membrane builder (8, 9). The composition was based on the asymmetric lipid distribution recently determined for red blood cells. It is featured by an increased phospholipid unsaturation in the cytoplasmic leaflet, a strong preference of phosphatidylcholine (PC) and sphingomyelin (SM) lipids within the exoplasmic leaflet, and PC, phosphatidylethanolamine (PE), and anionic phosphatidylserine (PS) lipids within the cytoplasmic leaflet (10). The detailed lipid composition is given in Table S1. To prevent inter-leaflet tension, simulations of symmetric lipid bilayers were performed, with the outer and inner leaflet composition, respectively. After an equilibration of 100 ns, the overall area per lipid was determined for each symmetric lipid bilayer (0.43 $\mathrm{nm}^2$ for the outer leaflet composition, 0.45 $\mathrm{nm}^2$ for the inner). The number of lipids were scaled according to the ratio of this values, to construct an asymmetric membrane with 8 x 8 $\mathrm{nm}^2$ lateral area.

The receptor-membrane systems were solvated in 8 x 8 x 13 $\mathrm{nm}^3$ rectangular boxes (11 x 11 x 15 $\mathrm{nm}^3$ for Fc$\gamma$RIIb$^{glyc}$) filled with a 0.15 M NaCl solution using the TIP3P water model (11, 12). All simulations were performed using the GROMACS 2020.1 and GROMACS 2021.6 packages (13–16) and the CHARMM36m force field for proteins and lipids (12, 17, 18). Parameters for the sugar residues were taken from the CHARMM36 forcefield for carbohydrates (19–21) and were attached to each potential N-glycosylation site of the human Fc$\gamma$RIIb using the CHARMM-GUI webserver (8, 9). For initial energy minimisation, 5,000 steps of the steepest descent algorithm were performed, followed by short equilibration runs with time steps of 1 fs and 2 fs. Production runs were performed at 310 K and 1 bar with a time step of $\Delta t = 2$ fs. Temperature and pressure coupling were achieved using the Nosé-Hoover thermostat (22, 23) (v-rescale thermostat (24) with $\tau_T = 0.1$ ps for Fc$\gamma$RIIb$^{glyc}$) and the semi-isotropic Parrinello-Rahman pressure coupling scheme (25) with coupling constants $\tau_T = 1.0$ ps and $\tau_P = 5.0$ ps, respectively. Periodic boundary conditions were applied for all simulations. Lennard-Jones interactions were shifted to zero between 1.0 nm and 1.2 nm. The particle-mesh Ewald method (26) with a cutoff of 1.2 nm was used to compute the (long-range) electrostatic interactions.

**Glycosylation.** Analysis for the Fc$\gamma$RIIb$^{glyc}$ was performed in the same way as described for the non-glycosylated receptors. Overall, the results were found to be hardly affected by glycosylation. Results for the inclination angle as a function of time and the distribution of sampled ectodomain configurations are provided in Figures S2 and S3. Biased by the shorter simulation times for Fc$\gamma$RIIb$^{glyc}$ of 10 µs (four replicas) compared to 20 µs for Fc$\gamma$RIIb$^{wt}$ and Fc$\gamma$RIIb$^{mut}$ (see also Figure S2), a larger fraction of configurations is accessible for IgG binding.

**Simulation analysis.** The tilt angle of the TM domain of Fc$\gamma$RIIb was defined as the angle between the helical axis (residues M222-R248) and the membrane normal ($z$-axis). The kink angle was defined as the angle between the center of mass (COM) of $C_\alpha$- and N-atoms of the first three TMD residues (S218, S219, S220), the COM of the $C_\alpha$- and N-atom of residue 232 (I232/T232) and the COM of $C_\alpha$- and N-atoms of

**Table S1. Lipid composition of the asymmetric bilayer. Listed are the numbers of respective lipids within the leaflets and their fraction on the total number of lipids within the leaflet (in brackets).**

| Lipid type | cytosolic leaflet | extracellular leaflet |
|---|---|---|
| CHOL | 59 (0.42) | 60 (0.40) |
| PLPC (16:0 / 18:2) | 13 (0.09) | 22 (0.15) |
| SOPC (18:0 / 18:1) | - | 10 (0.07) |
| PAPC (16:0 / 20:4) | - | 8 (0.05) |
| POPC (16:0 / 18:1) | 5 (0.04) | - |
| DPPC (16:0 / 16:0) | 3 (0.02) | - |
| POPE (16:0 / 18:1) | 3 (0.02) | - |
| PDoPE (16:0 / 22:6) | 11 (0.08) | - |
| SAPE (18:0 / 20:4) | 5 (0.04) | - |
| PAPS (16:0 / 20:4) | 19 (0.13) | - |
| SAPS (18:0 / 20:4) | 2 (0.01) | 2 (0.01) |
| PIP2 (18:0 / 20:4) | 3 (0.02) | - |
| PSM (18:1 / 16:0) | 2 (0.01) | 18 (0.12) |
| NSM (18:1 / 24:1) | - | 14 (0.09) |
| LSM (18:1 / 24:0) | - | 12 (0.08) |
| PEp (18:0 / 20:4) | 17 (0.12) | 4 (0.03) |

The number of lipids was scaled by 1.52 for the Fc$\gamma$RIIb$^{glyc}$ to maintain the molar lipid ratios in both leaflets. CHOL: Cholesterol; PLPC: 1-Palmitoyl-2-linoleoyl-sn-glycero-phosphatidylcholine; SOPC: 1-Stearoyl-2-oleoyl-sn-glycero-phosphatidylcholine; PAPC: 1-Palmitoyl-2-arachidonoyl-sn-glycero-phosphatidylcholine; POPC: 1-Palmitoyl-2-oleoyl-sn-glycero-phosphatidylcholine; DPPC: 1,2-Dipalmitoyl-sn-glycero-phosphatidylcholine; POPE: 1-Palmitoyl-2-oleoyl-sn-glycero-phosphatidylethanolamine; PDoPE: 1-Palmitoyl-2-docosahexaenoyl-sn-glycero-phosphatidylethanolamine; SAPE: 1-Stearoyl-2-arachidonoyl-sn-glycero-phosphatidylethanolamine; PAPS: 1-Palmitoyl-2-arachidonoyl-sn-glycero-phosphatidylserine; SAPS: 1-Stearoyl-2-arachidonoyl-sn-glycero-phosphatidylserine; PIP2: 1-Stearoyl-2-arachidonoyl-sn-glycero-phosphatidylinositol-3,5-bisphosphate; PSM: N-Palmitoyl-D-erythro-sphingosylphosphatidylcholine; NSM: N-Nervonoyl-D-erythro-sphingosylphosphatidylcholine; LSM: N-lignoceroyl-D-erythro-sphingosylphosphatidylcholine; PEp: 1-Octadecenyl-2-arachidonoyl-sn-glycero-phosphatidylethanolamine.

the last three TMD residues (R248, K249, K250). The inclination angle of the ectodomain was defined as the angle between the membrane normal (z-axis of simulation box) and a vector linking the COM of $C_\alpha$- and N-atoms of V161, N199, L204 and the COM of $C_\alpha$- and N-atoms of H188, E144, Q215.

To assess whether specific orientations of the Fc$\gamma$RIIb ectodomain with respect to the membrane are compatible for antibody binding, the IgG structure (PDB entry: 1IGY (27)) was aligned to the Fc$\gamma$RIIb ectodomain according to the binding position obtained from the IgG-Fc$\gamma$R crystal structure (PDB entry: 5VU0 (28)). If the position of any IgG atom was less than 7 Å above the mean position of the lipid headgroup nitrogen atoms in the extracellular leaflet, the Fc$\gamma$RIIb ectodomain orientation was considered incompatible for IgG binding.

For the diffusion coefficients of the lipids, the lateral mean squared displacement (MSD) was calculated, using the center of mass (COM) of the headgroup heavyatoms as reference, except for cholesterol were the hydroxyl oxygen was used. For the diffusion of the receptor, the COM of the TMD was used as reference. Diffusion coefficients were obtained by a least squared fit of the linear region (50-150 ns).

Lateral positions of the phosphorus atom in the phosphodiester linkage of each individual PIP2 lipid were followed in 10 ns time-steps and plotted with an *ad hoc* Python script.

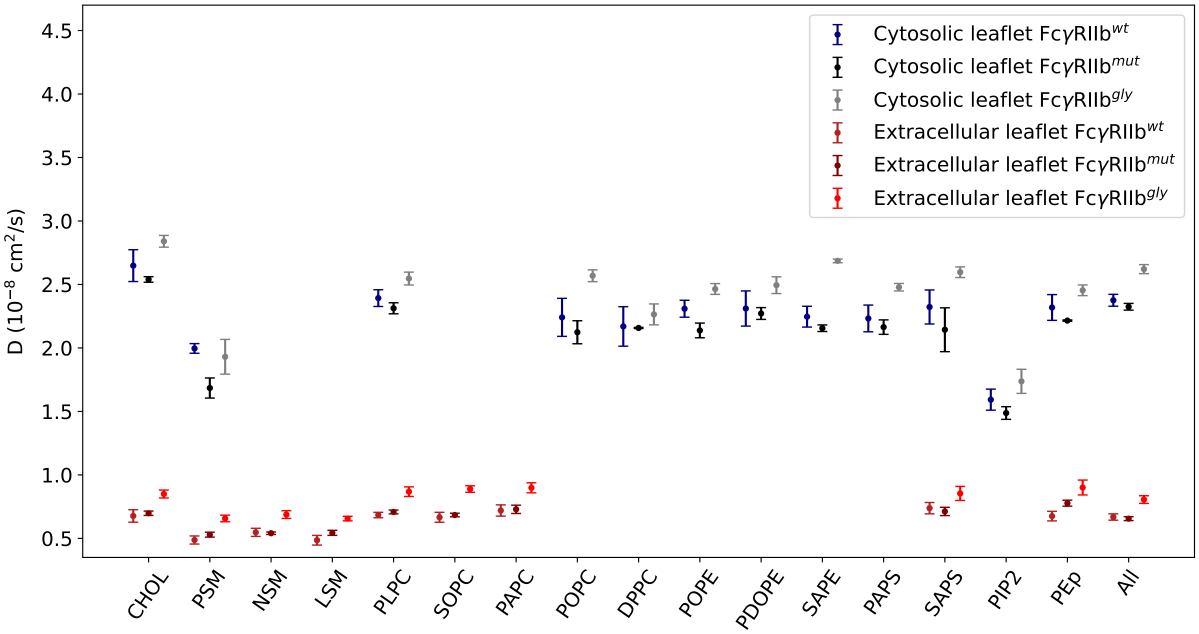


**Fig. S1.** Diffusion coefficients of all lipid types, separately for the cytosolic and the extracellular leaflets of the plasma membrane model. Error bars represent $\pm$ standard error of the mean (SEM).

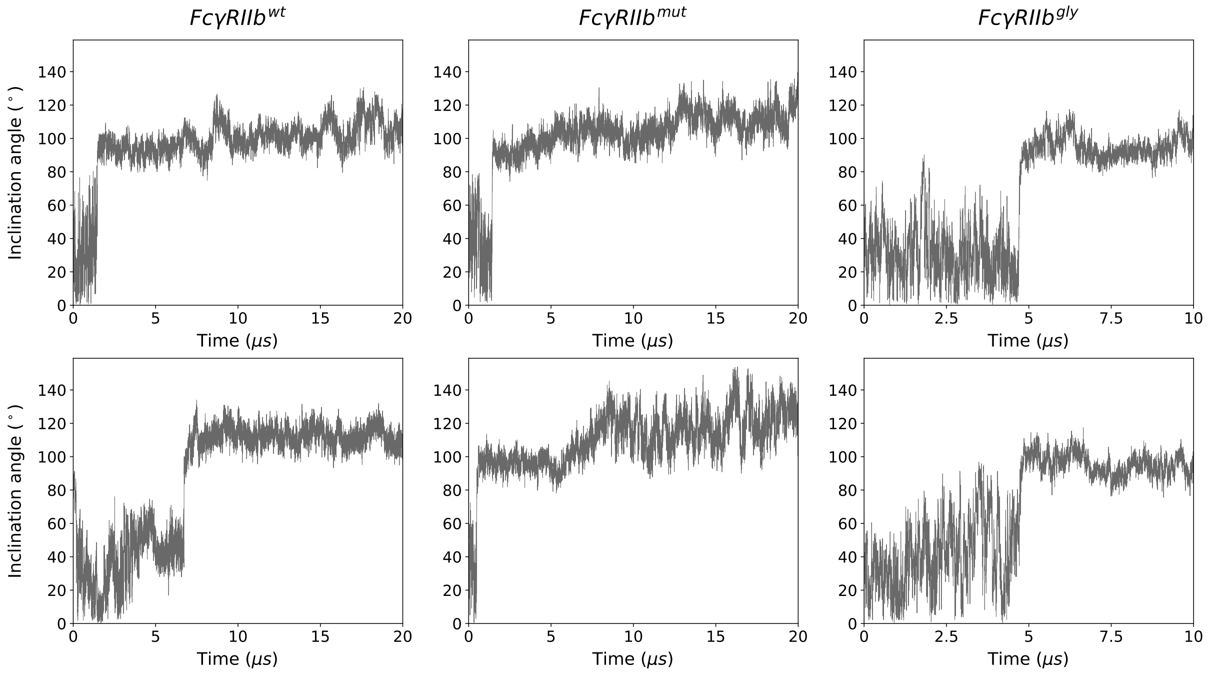


**Fig. S2.** Ectodomain inclination angle for two representative MD simulations of Fc$\gamma$RIIb wild type, the Fc$\gamma$RIIb-I232T mutant, and the glycosylated Fc$\gamma$RIIb, respectively, as a function of simulation time.

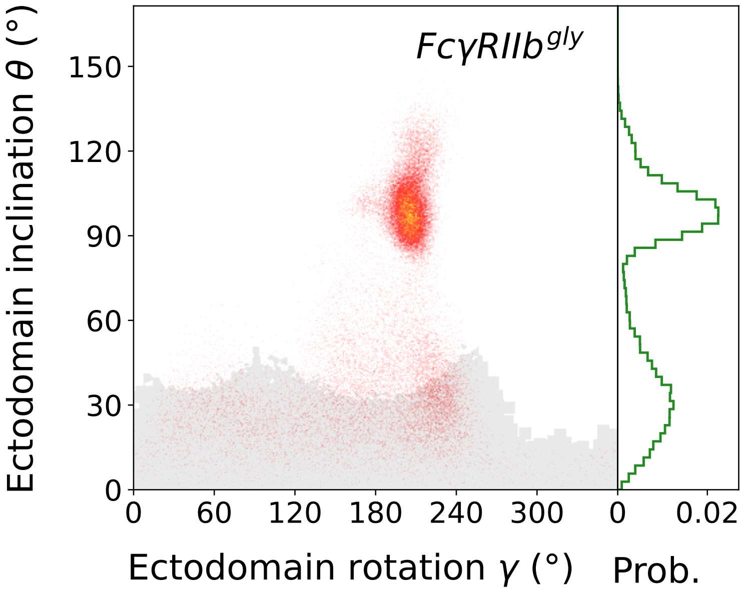


**Fig. S3.** Shown is the distribution of ectodomain configurations in the $\theta$-$\gamma$ space, including the data from four replicas, and the distribution of ectodomain inclination angles. Fc$\gamma$RIIb$^{glyc}$ ectodomain orientations within the *gray shaded* area were compatible with the binding of IgG (30.7 % of all observed conformations).

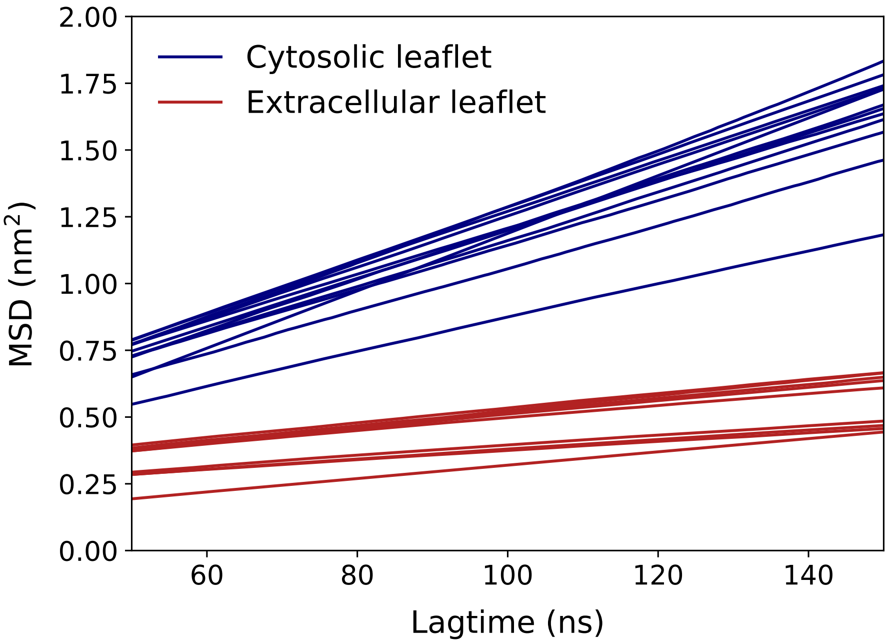


**Fig. S4.** Mean square displacement (msd) as a function of time. Each line corresponds to one lipid type within the respective leaflet (*red lines*: extracellular leaflet, *blue lines*: cytosolic leaflet).

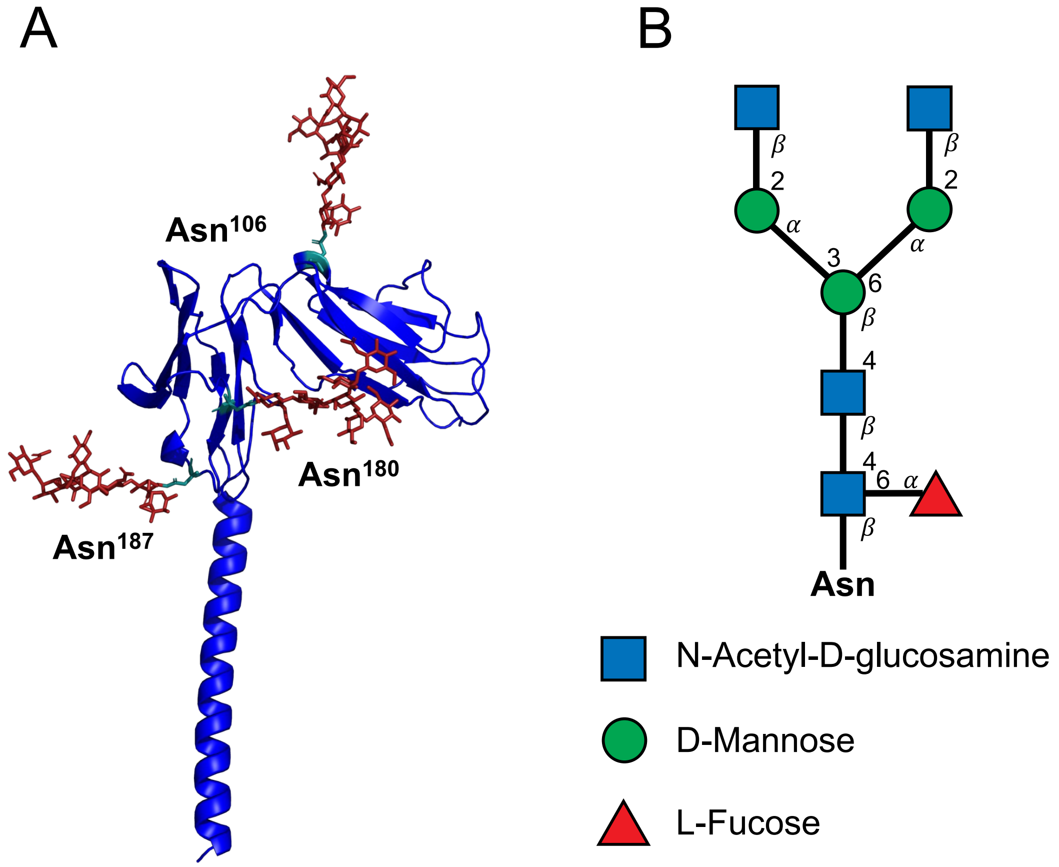


**Fig. S5.** Shown is Fc$\gamma$RIIb receptor with glycosylation sites within the ectodomain. (*A*) shows the N-glycosylated asparagine residues in the extracellular domain. The sugar residues with the linked amino acid are shown as sticks. The extracellular and TM domains of the receptor are shown in cartoon representation colored in blue. (*B*) Schematic structure of the added N-Glycan according to the Symbol Nomenclature for Glycans (SNFG) (29, 30).

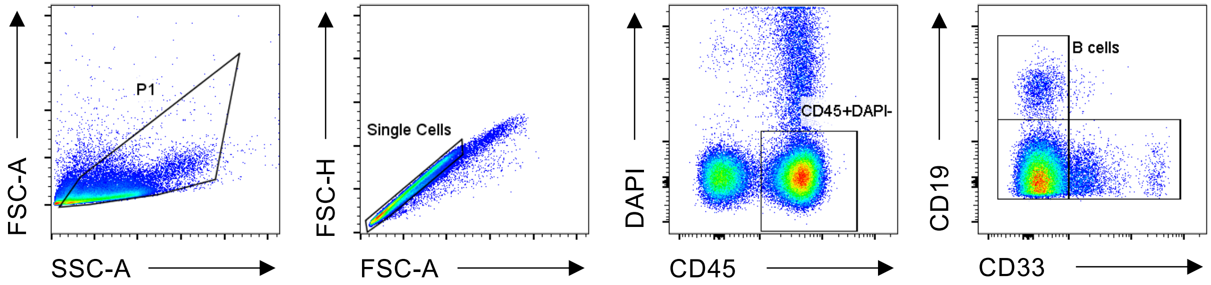


**Fig. S6.** Gating strategy of human peripheral blood leukocytes. Cells were determined based on forward and side scatter characteristics before doublet exclusion. B cells were identified as CD19+ CD33- cells within the living (DAPI-) CD45+ leukocytes. One exemplary sample is shown. Samples were acquired on a BD FACSCantoII.

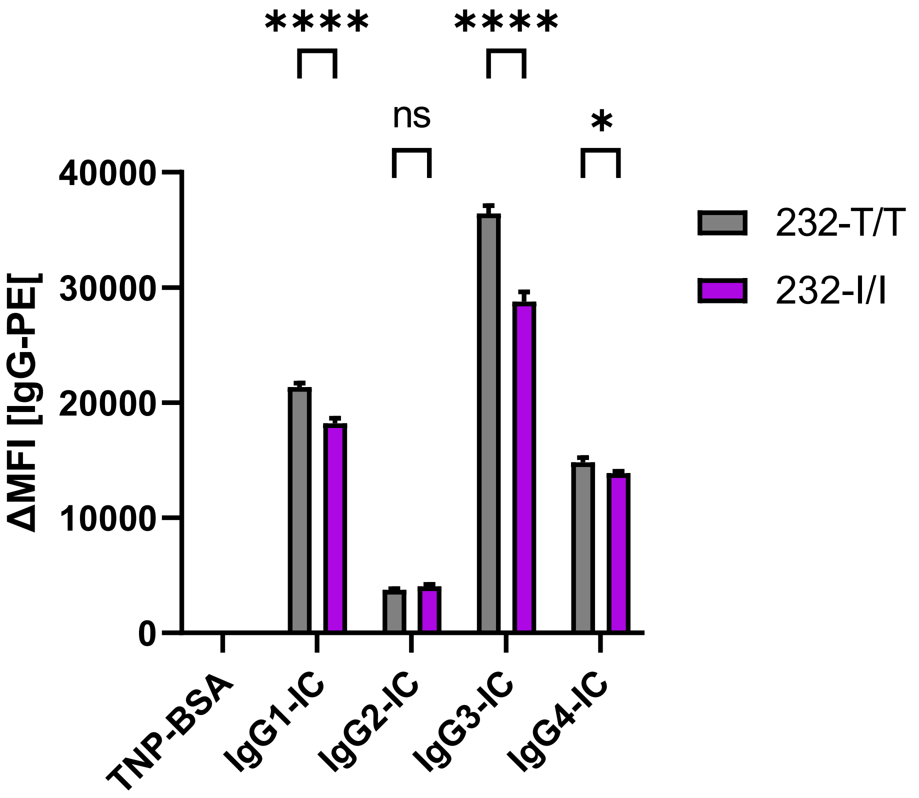


**Fig. S7.** Fc$\gamma$RIIb-I232T polymorphism does not affect IgG-IC binding. Interaction of IgG1-4 immune complexes with primary B cells (isolated from cord blood) homozygously expressing either Fc$\gamma$RIIb-232I or -232T was assessed by flow cytometry. Quantification of IgG-IC binding performed to 232I ($n = 3$, shown in violet) or 232T ($n = 3$, shown in dark gray) expressing B cells. Bars show mean and standard deviation of the median fluorescence intensity (MFI) upon detection of bound IgG-IC using fluorescently labeled anti-human IgG F(ab)2. TNP-BSA antigen is shown as negative control and background fluorescence of PBS-treated cell was substracted. Statistical analysis was performed by 2-way ANOVA. n.s. not significant, *p<0.05, **** p<0.0001.

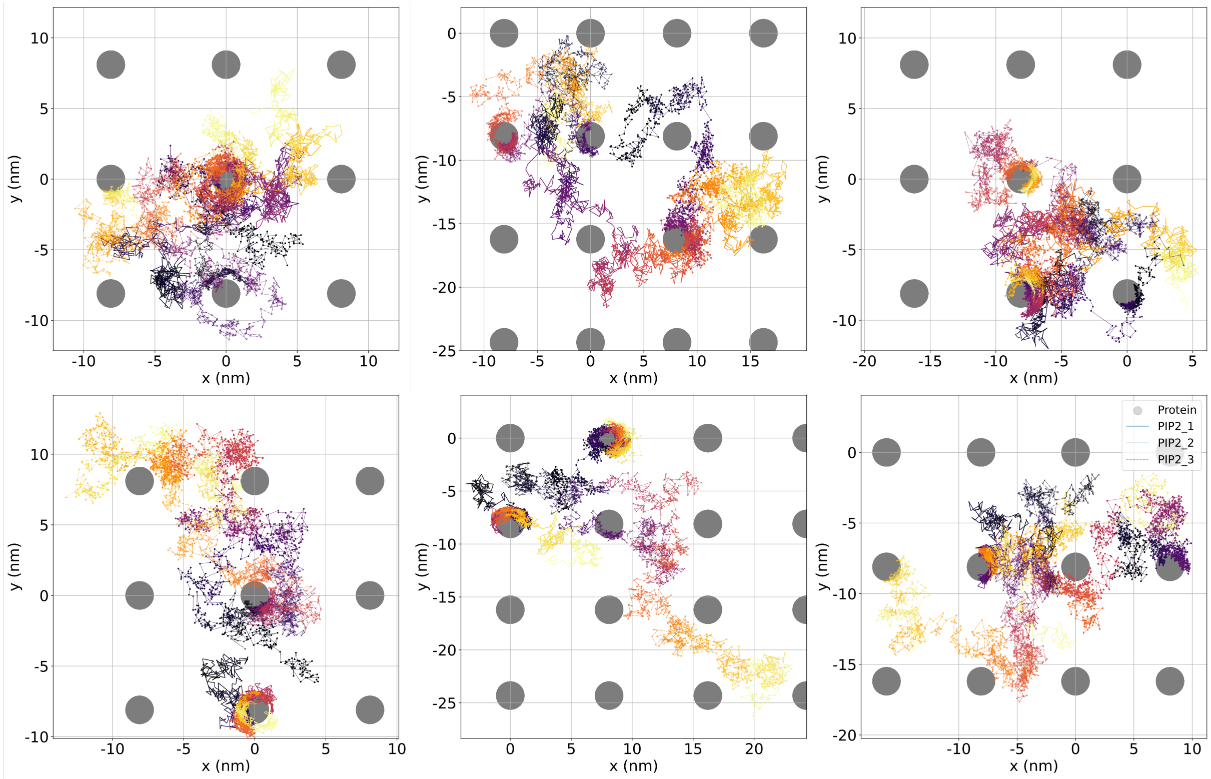


**Fig. S8.** Lateral membrane position of PIP2 molecules for selected simulations of the Fc$\gamma$IIb wt receptor embedded in a plasma membrane model. The area occupied by the protein is depicted as a filled gray circle and replicated for 8-11 periodic images. PIP2 movements are shown as lines colored with gradient from dark purple to light yellow, tracking from $t = 0\mu$s to $t = 20\mu$s.

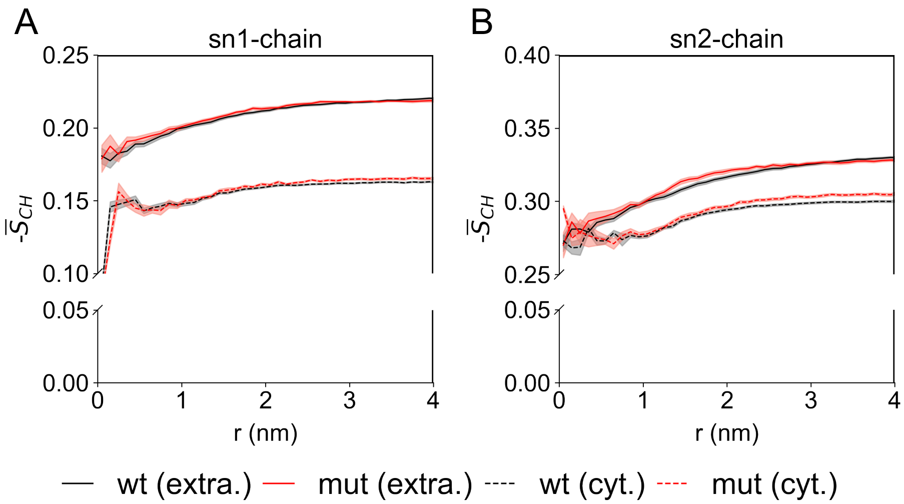


**Fig. S9.** Mean lipid order parameters, averaged over the first 14 CH2 groups of each acyl chain. The averages were binned as a function of distance $r(nm)$ between the lipid headgroups (not distinguishing between the different lipid types) and the Fc$\gamma$IIb TM domain. The order is shown for lipids of the extracellular leaflet ('extra.') and lipids of the cytosolic leaflet ('cyt.'). Data were averaged over the final 10 $\mu$s of wt and I232T mutant simulations, with the shaded area representing the standard error of the mean calculated over all replica simulations.

### References


1. Sondermann P, Huber R, Jacob U (1999) Crystal structure of the soluble form of the human Fcgamma-receptor IIb: a new member of the immunoglobulin superfamily at 1.7 A resolution. *EMBO J.* 18(5):1095–1103.
2. Webb B, Sali A (2016) Comparative protein structure modeling using MODELLER. *Curr Protoc Bioinformatics* 54(1):5–6.
3. Hayes JM, et al. (2014) Fc gamma receptor glycosylation modulates the binding of IgG glycoforms: a requirement for stable antibody interactions. *J. Proteome Res.* 13(12):5471–5485.
4. Hayes JM, et al. (2017) Identification of Fc gamma receptor glycoforms that produce differential binding kinetics for Rituximab. *Mol. Cell. Proteomics* 16(10):1770–1788.
5. Cosgrave EFJ, et al. (2013) N-linked glycan structures of the human Fc$\gamma$ receptors produced in NS0 cells. *J. Proteome Res.* 12(8):3721–3737.
6. The UniProt Consortium (2023) UniProt: the Universal Protein Knowledgebase in 2023. *Nucleic Acis Research* 51:D523–D531.
7. Varki A, et al., eds. (2022) *Essentials of Glycobiology.* (Cold Spring Harbor Laboratory Press, Cold Spring Harbor (NY)).
8. Jo S, Kim T, Iyer V, Im W (2008) CHARMM-GUI: a web-based graphical user interface for CHARMM. *J. Comput. Chem* 29(11):1859–1865.
9. Lee J, Cheng X, Swails J, Yeom M, Eastman Pea (2016) CHARMM-GUI input generator for NAMD, GROMACS, AMBER, OpenMM, and CHARMM/OpenMM simulations using the CHARMM36 additive force field. *J. Chem. Theory Comput.* 12(1):405–413.
10. Lorent J, et al. (2020) Plasma membranes are asymmetric in lipid unsaturation, packing and protein shape. *Nat. Chem. Biol* 16(6):644–652.
11. Jorgensen WL, Chandrasekhar J, Madura JD, Impey RW, Klein ML (1983) Comparison of simple potential functions for simulating liquid water. *J. Chem. Phys.* 79(2):926–935.
12. Yu Y, Klauda JB (2020) Update of the CHARMM36 united atom chain model for hydrocarbons and phospholipids. *J. Phys. Chem. B* 124(31):6797–6812.
13. Van Der Spoel D, et al. (2005) GROMACS: fast, flexible, and free. *J. Comput. Chem* 26(16):1701–1718.
14. Abraham M, et al. (2015) GROMACS: High performance molecular simulations through multi-level parallelism from laptops to supercomputers. *SoftwareX* 1:19–25.
15. Lindahl E, Abraham M, Hess B, van der Spoel D (2020) GROMACS 2020.1 Source code.
16. Lindahl E, Abraham M, Hess B, van der Spoel D (2020) GROMACS 2021.6 Source code.
17. Klauda JB, et al. (2010) Update of the CHARMM all-atom additive force field for lipids: validation on six lipid types. *J. Phys. Chem. B* 114(23):7830–7843.
18. Huang J, et al. (2017) CHARMM36m: an improved force field for folded and intrinsically disordered proteins. *Nat. Methods* 14(1):71–73.
19. Guvench O, et al. (2008) Additive empirical force field for hexopyranose monosaccharides. *J. Comput. Chem.* 29(15):2543–2564.
20. Guvench O, Hatcher ER, Venable RM, Pastor RW, Mackerell AD (2009) CHARMM additive All-Atom force field for glycosidic linkages between hexopyranoses. *J. Chem. Theory Comput.* 5(9):2353–2370.
21. Guvench O, et al. (2011) CHARMM additive all-atom force field for carbohydrate derivatives and its utility in polysaccharide and carbohydrate-protein modeling. *J. Chem. Theory Comput.* 7(10):3162–3180.
22. Nosé S (1984) A unified formulation of the constant temperature molecular dynamics methods. *J. Chem. Phys.* 81(1):511–519.
23. Hoover WG (1985) Canonical dynamics: Equilibrium phase-space distributions. *Phys. Rev. A* 31(3):1695.
24. Bussi G, Donadio D, Parrinello M (2007) Canonical sampling through velocity rescaling. *J. Chem. Phys.* 126(1):014101.

25. Parrinello M, Rahman A (1981) Polymorphic transitions in single crystals: A new molecular dynamics method. *J. Appl. Phys.* 52(12):7182–7190.
26. Darden T, York D, Pedersen L (1993) Particle mesh Ewald: An Nlog(N) method for Ewald sums in large systems. *J. Chem. Phys.* 98(12):10089–10092.
27. Harris L, Skaletsky E, McPherson A (1998) Crystallographic structure of an intact IgG1 monoclonal antibody. *J. Mol. Biol.* 275(5):861–872.
28. Falconer D, Subedi G, Marcella A, Barb A (2018) Antibody fucosylation lowers the Fc$\gamma$RIIIa/CD16a affinity by limiting the conformations sampled by the N162-glycan. *ACS Chem. Biol* 13(8):2179–2189.
29. Varki A, et al. (2015) Symbol nomenclature for graphical representations of glycans. *Glycobiology* 25(12):1323–1324.
30. Neelamegham S, et al. (2019) Updates to the symbol nomenclature for glycans guidelines. *Glycobiology* 29(9):620–624.
